# Epithelial Outgrowth Through Mesenchymal Rings Drives Alveologenesis

**DOI:** 10.1101/2022.10.06.511160

**Authors:** Nicholas M. Negretti, Yeongseo Son, Philip Crooke, Erin J. Plosa, John T. Benjamin, Christopher S. Jetter, Claire Bunn, Nicholas Mignemi, John Marini, Alice N. Hackett, Meaghan Ransom, David Nichols, Susan H. Guttentag, Heather H. Pua, Timothy S. Blackwell, William Zacharias, David B. Frank, John A. Kozub, Anita Mahadevan-Jansen, Jonathan A. Kropski, Christopher V.E. Wright, Bryan Millis, Jennifer M. S. Sucre

## Abstract

Determining how alveoli are formed and maintained is critical to understanding lung organogenesis and regeneration after injury. While technological barriers have heretofore limited real-time observation of alveologenesis, we have now used scanned oblique plane illumination microscopy of living lung slices to observe specific cellular behaviors at high resolution over several days. Contrary to the prevailing paradigm that alveoli form by airspace subdivision via ingrowing septa, we find that alveoli form by ballooning epithelial outgrowth supported by stable mesenchymal ring structures. Our systematic analysis allowed creation of a computational model of finely-timed cellular structural changes that drive alveologenesis under normal conditions or with perturbed intercellular Wnt signaling. This new paradigm and platform can be leveraged for mechanistic studies and screening for therapies to promote lung regeneration.

**One-Sentence Summary:** Long-term live analysis of neonatal lungs supports a dynamic epithelial outgrowth model for alveologenesis.

---

Lung organogenesis results from precisely coordinated intercellular signaling pathways that induce cell-type specialization, and structural changes to generate the vast surface area required for gas exchange. The critical alveolar stage of lung development continues through early childhood in humans, equivalent to postnatal days (P) 5-28 in mice. Most previous explorations of alveolar development and structure have relied on either inferences from staged 2D (5-10 μm) histological sections or relatively short-term live imaging (<24 hrs) that was limited because of photobleaching and phototoxicity (*1*). We circumvented these hurdles using a tailored version of scanned oblique-plane illumination (SOPi) microscopy (*2*) to perform long-term (72 hrs), high-resolution live analysis of alveologenesis in precision-cut lung slices (PCLS) *ex vivo*. Using this 4-dimensional (4D) imaging platform with PCLS from transgenic reporter mice we have tracked the reproducible behaviors of epithelial and mesenchymal cells in the development of new alveoli.

Vibratome-sectioned PCLS (300-500 μm thick) from P5 neonatal mice at the saccular to alveolar transition **(Fig. 1A)** provided an *ex vivo* model of alveologenesis, that followed the *in vivo* process with good fidelity. PCLS continued to develop and form new alveolar structures **(Fig. 1B)** (*3, 4*). Of note, PCLS contain all relevant cell types of lung parenchyma, including those producing matrix and growth factors to support growth in medium free of additives (*5*). Comparing adjacent PCLS from the same lung fixed immediately at P5 to those cultured for 48 hrs revealed significant changes in airspace volume over time (**Fig. 1C)** and increased alveolar surface area. P5 PCLS cultured for 48 hrs showed tissue structures essentially highly similar to lungs taken directly from P7 mice (**Fig. 1D, E)**.

**Fig. 1.**
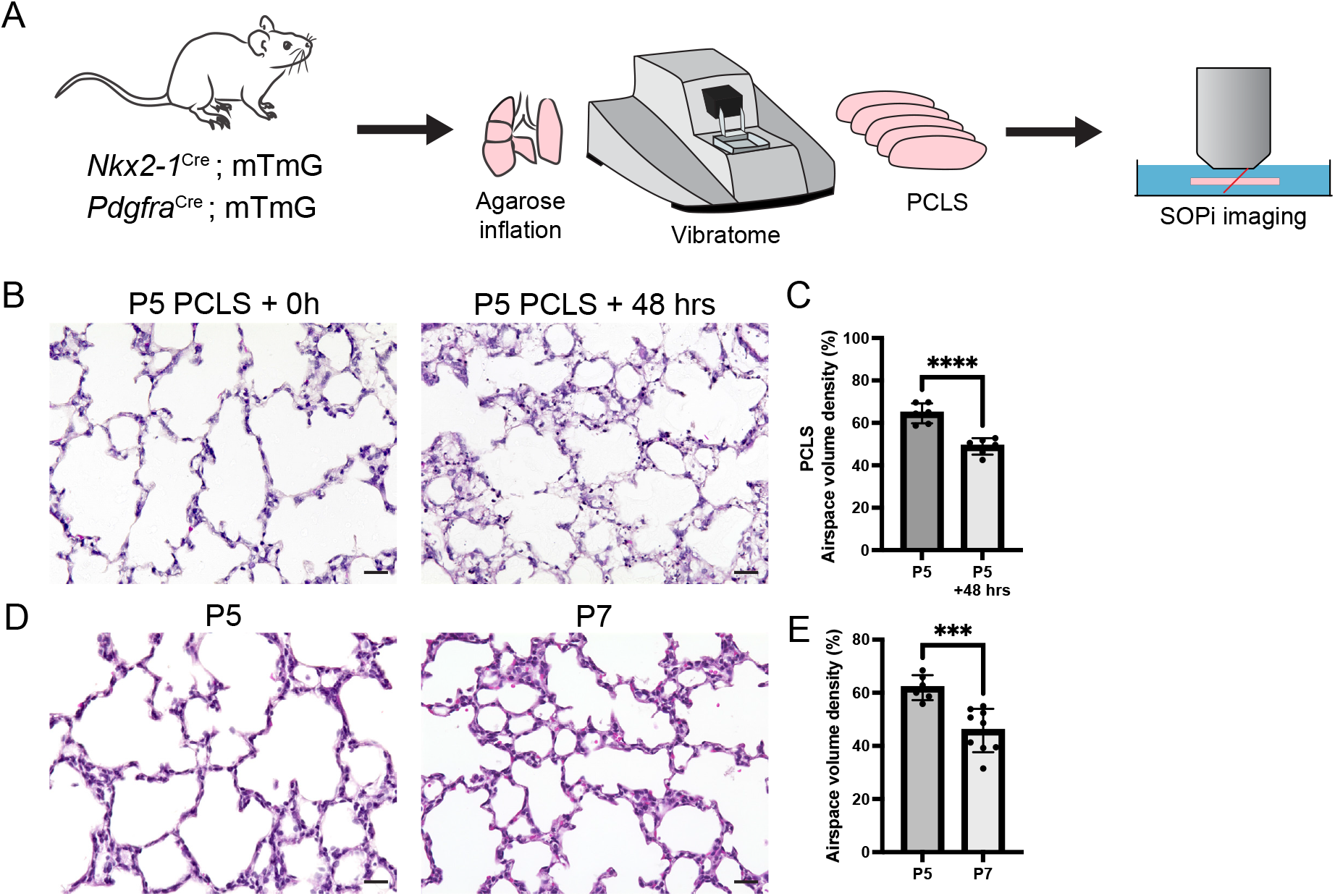
Precision-cut lung slices (PCLS) model alveologenesis *ex vivo*. **(A)** Five-day-old mT/mG transgenic mice fluorescently reporting for alveolar epithelial (Nkx2-1-Cre) or alveolar mesenchymal cells (Pdgfra-Cre) were imaged by scanned oblique-plane illumination (SOPi) microscopy. **(B)** PCLS were fixed immediately after preparation at P5, or after 48 hrs in culture and hematoxylin/eosin (H&E) stained. **(C)** Airspace volume density was calculated and compared from PCLS created from the same lung at P5 and P5 + 48 hours (six mice from two litters), with structural changes equivalent in many respects to **(D)** the H&E sections from P5 or P7 mice and quantification of airspace volume density (**E)**; *\*\*\*p*<0.001, *\*\*\*\*p<0*.0001.

To label and follow epithelial or mesenchymal cells during early alveologenesis, we used mT/mG Cre-switched transgenic mice that express membrane-bound tdTomato (mT) before Cre-excision but membrane-targeted green fluorescent protein (mG) after excision (*6*). PCLS from mice reporting for epithelial cells (Nkx2-1-Cre (*7*)) or alveolar mesenchymal cells (Pdgfra-Cre (*8*)) were informatively imaged over a depth of 100-150 μm. When viewed in 3D, P5 PCLS from mT/mG;Pdgfra-Cre mice contained an extensive network of connected ring-like structures, each formed by four to five Pdgfra-traced cells (**Fig. 2A)**. This observation is consistent with the fishnet-like mesenchymal web previously proposed to support alveolar development (*9*). We analyzed single optical planes from several individual *z*-stacks, finding that any Pdgfra+-traced alveolar fibroblasts that appeared in 2D as putative “tips of in-growing septa” (**Fig. 2B)** were always, in fact, part of the internal ring structures. Similarly, 3D imaging of Nkx2-1-traced cells allowed a more precise derivation of alveolar architecture (**Fig. 2C**) than 2D (**Fig. 2D**). Notably, epithelial cells that seemed to be part of an ingrowing septal wall when viewed in a single plane were actually an extended surface of the ballooning alveolus oriented perpendicular to the *x-y* optical sectioning plane. Cells expressing markers of alveolar type 2 (AT2) cells (*Sftpc*) and alveolar type 1 (AT1) cells (*Hopx*) are present in our *ex vivo* model (**Fig S1A)**, and the rounded or highly flattened Nkx2-1-traced cells fit the characteristics of AT2 cells and AT1 cells, respectively.

**Fig. 2.**
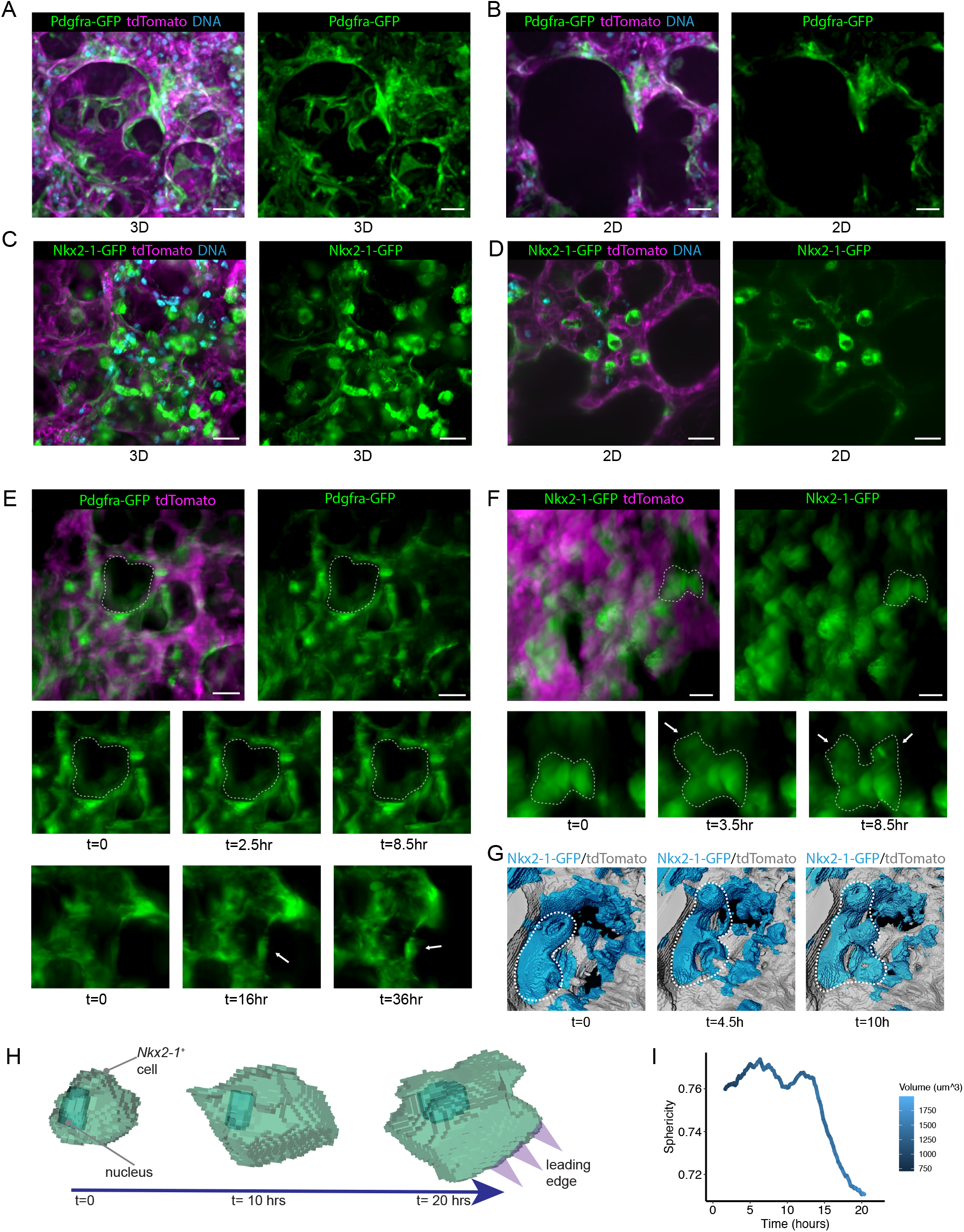
Alveologenesis proceeds by epithelial outgrowth from a mesenchymal support ring. **(A)** PCLS from mT/mG;Pdgfra-Cre mice were volumetrically imaged and displayed as a 3D projection. Left: GFP green, tdTomato magenta, DNA cyan. Right: GFP channel alone. **(B)** 2D projection from a single plane of the *z*-stack in A. **(C)** mT/mG;Nkx2-1-Cre PCLS similarly imaged and 3D projected. Left: GFP green, tdTomato magenta, DNA cyan. Right: GFP channel alone. **(D)** 2D single-plane projection from the *z*-stack in B. **(E)** mT/mG;Pdgfra-Cre PCLS imaged in 3D over time. White dotted outline: a single mesenchymal ring. **(F)** mT/mG;Nkx2-1-Cre PCLS mice imaged in 3D over time. White outline: an epithelial cluster. **(G)** Surface rendering of lung tissue from mT/mG;Nkx2-1-Cre PCLS. GFP+ cells in cyan, all other cells gray. **(H)** A representative single Nkx2-1-tracked GFP+ cell tracked over time and surface rendered with nucleus indicated. **(I)** Sphericity and cellular volume of representative flattening epithelial cell over time. Scale bars = 25 μm.

Most Pdgfra-traced ring structures were present when imaging started with additional rings forming over 72-hrs of imaging (**Fig. 2E, Supplementary Video 1)**. Multiple measurements of ring size over time revealed no significant change in diameter (**Fig S1B)**. Nkx2-1-traced epithelial cells clustered adjacent to the ring **(Fig S2**), followed by directional extrusion through the ring, extension, and flattening, collectively undergoing an outward ballooning movement (**Fig. 2F, G, Supplementary Video 2)**. With extrusion, the soma of each cell became displaced away from the ring approximately 60-100 μm, approximating the normal alveolar diameter in P5 mice (*10*). Without any apparent difference across the 3.4 million μm^3^ of each volume imaged over multiple PCLS, we observed a reproducible sequence of epithelial cell clustering and extrusion, followed by elongation and flattening **(Fig. 2G, S3A Supplementary Video 2, 3)**. Alveologenesis events were broadly asynchronous across any single PCLS (**Supplementary Video 2)**. Notably, alveoli formed in single-extrusion and adjacent dual-extrusion structures, and rarely as conjoined groups of three-to-five alveologenesis events (**Fig. 2G, Supplementary Video 2, 3)**. This behavior was expected because a characteristic feature of lung development is the temporally concurrent formation of multiple adjoined alveoli with common separating walls. Despite extensive 4D imaging, we found no instances representing dynamically ingrowing epithelial/mesenchymal septal walls that could drive alveologenesis by airspace subdivision. We conclude that such processes are rare or non-existent, and that ring-proximal epithelial cell clustering followed by directional extrusion and cell flattening (“ballooning”), as cells transition to mature AT1 cells, represents the predominant alveologenesis mechanism.

As mentioned above, AT1 differentiation involves large-scale cell-shape alteration – from rounded to extremely thin and outspread – for these gracile cells to assume gas exchange function. Tracking mG fluorescence in Nkx2-1-traced cells allowed cell-shape determination over time **(Fig 2H)**. By a sphericity metric of relative roundness/flatness, we found that many cells underwent significant flattening during alveolar-structure formation, with ~5.2% of Nkx2-1-traced cells undergoing a complete, fairly rapid round-to-flat transition (**Fig S3D)**. After a variable lag period for each neo-alveolus imaged in this way (representing the aforementioned asynchronicity across the parenchyma), the flattening period was ~10 hrs (**Fig. 2I**). Our observations suggest a similar cellular rearrangement occurs in alveologenesis as reported in prior studies of the earlier saccular stage (E15.5-18.5), which found that epithelial-cell protrusion drives differentiation in explants (*11*). The epithelial clustering, outward extrusion and flattening occur with relative lack of influence from cell proliferation. Nuclear tracking failed to detect evidence of significant cell division during these events, and about ~1% of Nkx2-1-traced epithelial cells were Ki67^+^ (**Fig. S3B, C)**, consistent with very few proliferating AT1 or AT2 cells at P5 in transcriptomic atlases of the developing lung (*12*).

Taken together, our 4D imaging of later, postnatal lung development has newly revealed that alveologenesis is driven by epithelial collection at interconnected mesenchymal ring-support structures, with directional epithelial extrusion through the rings tightly accompanied by spreading and flattening to become mature AT1 cells. This step forward in understanding alveologenesis should afford a substantial shift in the biological questions to be pursued in which the idea that septal ingrowth creates alveoli is left behind. Instead, we propose investigating the mechanisms by which the stabilization and disappearance of the mesenchymal ring (*13*) acts a transient scaffold of the final parenchymal structure and whether or not the ~75 μm displacement distance somehow sets the initial neo-alveolar diameter.

A key feature of this SOPi system is a water-immersion objective allowing application of small-molecule pathway modulators onto the live tissue. To determine if such 4D imaging can efficaciously reveal normal and abnormal developmental processes, we imaged PCLS from Nkx2-1-Cre;mT/mG mice treated with modulators of Wnt signaling, a pathway essential for alveologenesis (*14*). PCLS were cultured with CHIR (potent activator of canonical Wnt signaling) or XAV-939 (potent pan-Wnt inhibitor by inhibition of tankyrase). With both perturbations, PCLS had significantly fewer alveologenesis events than controls (**Fig. 3A-C)**. With CHIR, some epithelial cells began to elongate, but then reverted to their original shape and position (**Fig. 3A, Supplementary Video 4)**. Using nuclear tracking as a proxy for cellular displacement, CHIR caused epithelial cells to move faster than controls (**Fig. 3D**), and we captured cells displaying oscillatory back-and-forth behavior while failing to complete extrusion and ballooning movements. By contrast, global Wnt inhibition caused epithelial cells to exhibit little to no movement (**Fig. 3C, Supplementary Video 5)**. The absence of directional processivity of nuclear movements with Wnt activation (**Fig. 3D-F)** was consistent across all our 4D imaging data. Such perturbation of an intercellular signaling pathway essential to normal lung development sheds light on the profound impacts on tissue rearrangement in normal alveologenesis and how disrupted signaling that might happen upon injury.

**Fig. 3.**
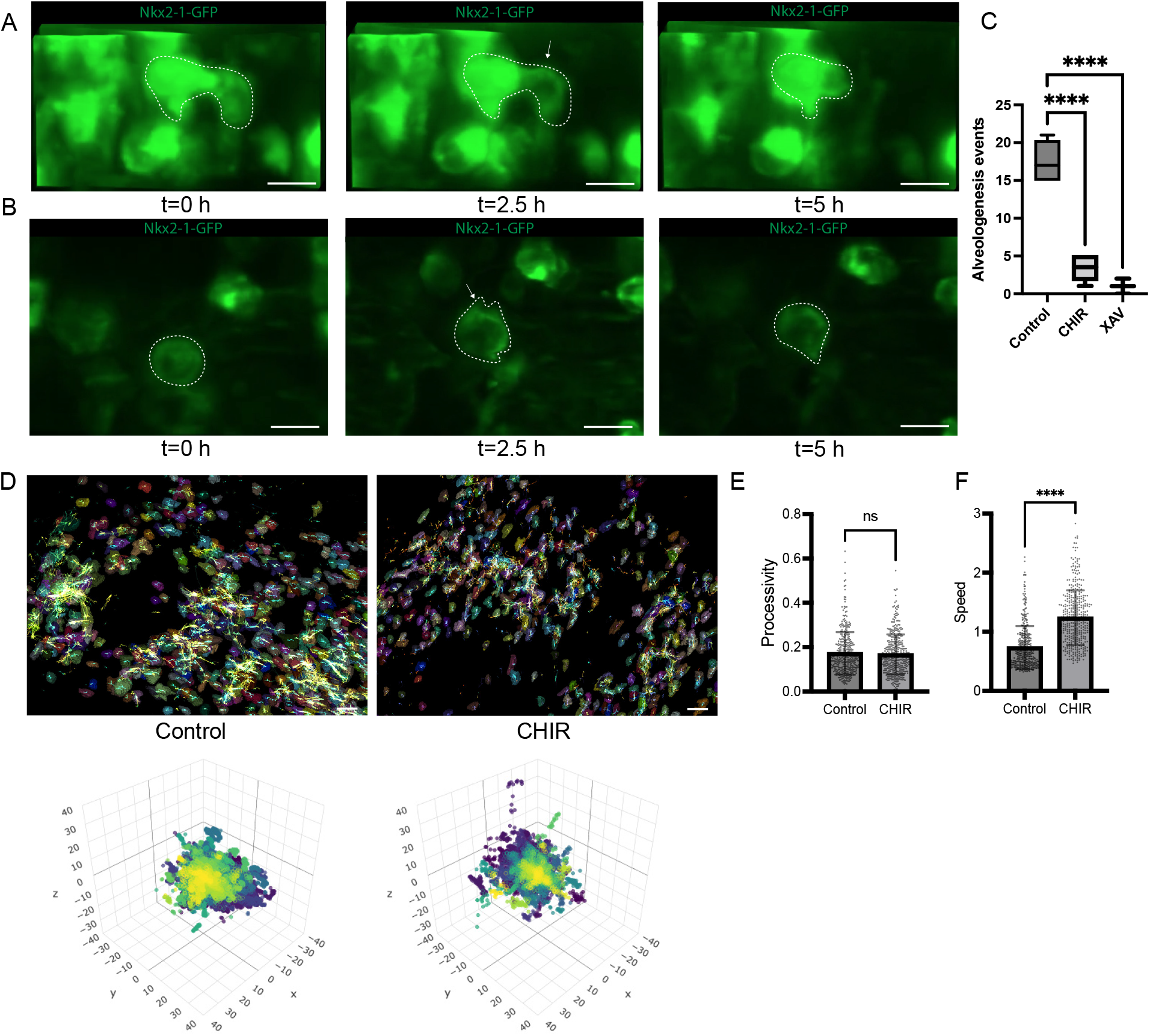
Alveologenesis is disrupted by perturbation of Wnt signaling. 3D projections from **(A)** CHIR or **(B)** XAV treated PCLS imaged over time. An individual epithelial cell is tracked over time (white outline). **(C)** Individual alveologenesis events by epithelial cells were scored in a blinded manner from PCLS from mT/mG;Nkx2-1-Cre tissue treated with CHIR99021 (Wnt activator), XAV 939 (Wnt inhibitor), or vehicle. \*\*\*\**p*<0.001 by one-way ANOVA. **(D)** Nuclei from control and CHIR-treated samples were tracked over time and cell trails (top) plotted in addition to Windrose plots (bottom). **(E)** Processivity (a measure of directed cell motion) and speed **(F)** was calculated from the nuclear motion. (**** p < 0.01 by Student’s *t*-test).

Alveolar mesenchymal cells are important sources of Wnt ligands during early alveologenesis, with precise spatiotemporal patterning of expression of *Wnt2* and *Wnt5a* (*12*). We speculate that Wnt inhibition prevented alveolar epithelial cell migration and subsequent extrusion, preventing even early stages of outgrowth-ballooning. Alternatively, it is possible that CHIR-based global Wnt activation may have overwhelmed the mesenchymal ring-sourced Wnt signaling gradient and disrupted the directional movement away from the Wnt source that allows full AT1 outspreading and maturation. Hence, our plausible model for normal stepwise structural development of alveoli is that localized, highly threshold-dependent Wnt signaling sourced from the mesenchymal ring initiates directional epithelial extrusion, and distance-dependent escape from Wnt signaling allows continued outspreading and flattening. In this model, displacement of cells away from ring-sourced Wnt to regions of lower ligand concentration allows a spatiotemporally appropriate rise to dominance of other signaling pathways to orchestrate progressive AT1 differentiation, consistent with waves of Wnt signaling being required for alveologenesis (*14*).

To quantify normal alveolar growth, we constructed a computational model from our membrane/cell shape-tracking data. Abstracting 3D fluorescence data from multiple alveoli into a computational matrix (**Fig. 4A-C)** allowed accurate modeling of the expanding epithelial perimeter and alveolar area over time **(Fig. 4A-C)**. Building and testing this model over multiple alveologenesis events fit well with the structural changes of epithelial outgrowth and alveolar expansion (**Fig. 4D)**. None of our observed alveologenesis events could be fitted to a septal-ingrowth paradigm. PCLS treated with CHIR or XAV-939 showed very little perimeter expansion compared to controls, consistent with the observations above that up- or down-regulating Wnt signaling greatly disturbs epithelial extrusion and alveolar growth. Such parameterization of alveologenesis should aid future rigorous comparisons between treatment conditions, genetic models, and developmental stages.

**Fig. 4.**
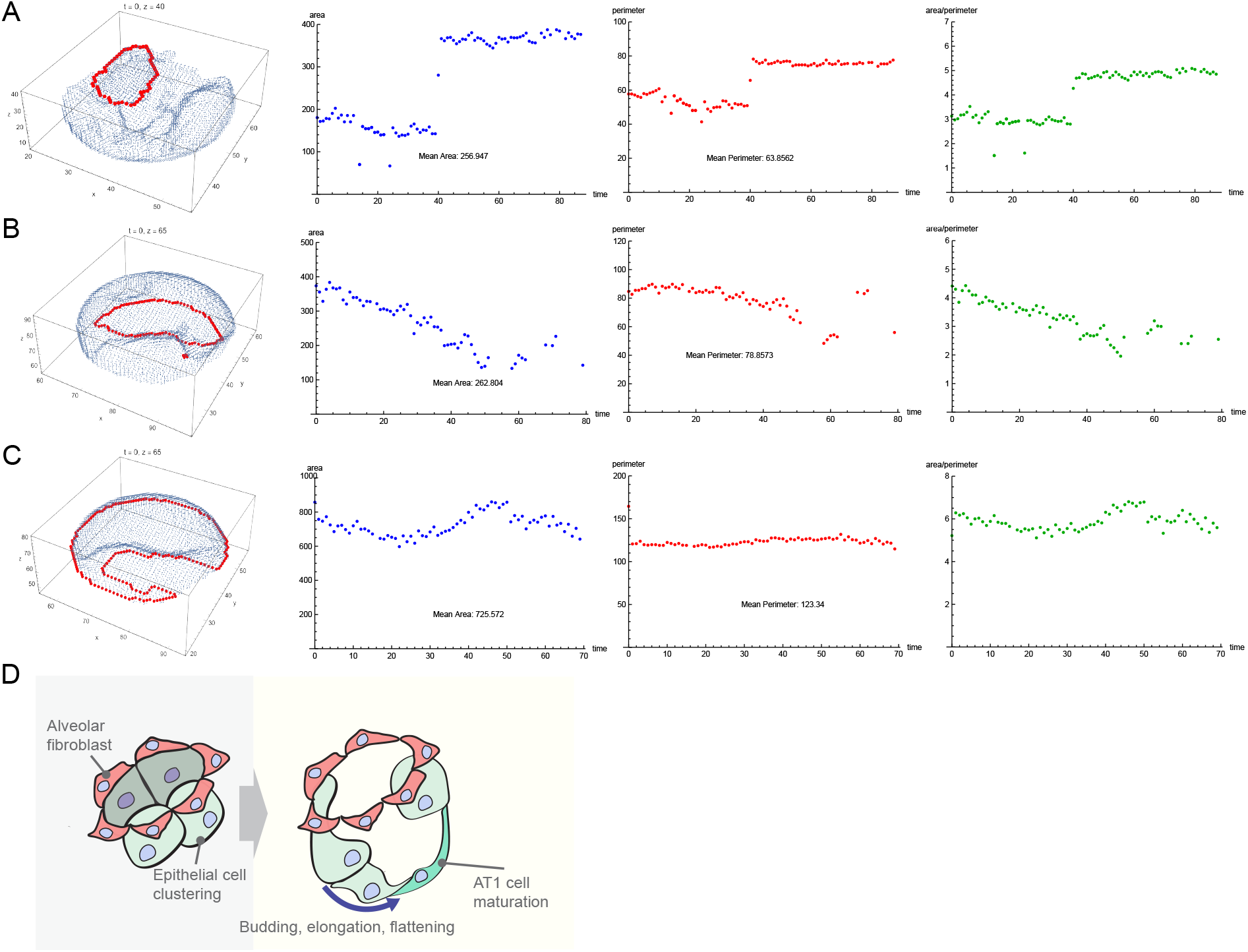
Computational modeling of alveologenesis based on membrane/cell shape-tracking data parameterizes epithelial outgrowth. (**A-C)** Computational modeling of representative single alveolar buds from **(A)** control, **(B)** CHIR-treated (Wnt activator) and **(C)** XAV-939 (Wnt inhibitor) treated PCLS imaged over time. Area, perimeter, and area/perimeter (a proxy of complexity) were calculated. **(D)** Proposed cartoon of alveologenesis characterized by epithelial outgrowth from a foundational mesenchymal ring.

These results rely on observations of lung slices *ex vivo* and submerged in liquid, therefore without an air interface. This feature of the experimental system probably does not present a substantial caveat to our conclusions because human alveologenesis begins *in utero* at ~32 wks gestation, independent of air inflation. Our system lacks possible modulatory effects of mechanical pressure caused by amniotic fluid and *in utero* breathing movements as well as a connected vasculature with a circulatory system. Nonetheless, the histological changes being consistent with expansion of surface area observed *in vivo* (*15*) strongly buttresses this *ex vivo* system as a relevant alveologenesis model.

In summary, this 4D *ex vivo* imaging system is well positioned not only to advance our understanding of the molecular-cellular mechanisms of alveologenesis, but also for evaluating effects of developmental lung injury on alveologenesis, and as a platform for drug discovery. Computational modeling of the timing and geometry of alveologenesis will direct us to finding how outward epithelial growth requires precisely organized epithelial-mesenchymal cellular and biomechanical interactions to drive this essential and sensitive process.

## Supplementary Materials

Materials and Methods

Figs. S1–S3

Movies S1 to S5

## METHODS

### Fluorescently labeled precision cut lung slices

The C57BL/6J background was used for all mouse experiments. For fluorescent imaging, Nkx2-1 Cre (Jackson Laboratories, Stock no.: 008661), or Pdgfra-Cre (Jackson Laboratories, Stock no.: 013148) mice were crossed with mT/mG mice (Jackson Laboratories, Stock no.: 007676) to generate Nkx-Cre;mTmG or Pdgfra-Cre;mTmG. The presence of the Pdgfra-Cre was tested with the following primer set: Pdgfra-Cre Forward: 5’-TCA GCC TTA AGC TGG GAC AT-3’, Pdgfra-Cre Reverse: 5’-ATG TTT AGC TGG CCC AAA TG-3’. The presence of Nkx2-1-Cre was tested with the following primer set: Nkx-Cre Forward: 5’-CTC TGG TGG CTG CCT AAA AC-3’, Nkx-Cre Reverse: 5’-CGG TTA TTC AAC TTG CAC CA-3’. The presence of the mT/mG allele was tested with the following primer set: Wild-Type Forward: 5’-CCG AAA ATC TGT GGG AAG TC-3’, Mutant Forward: 5’-CGG GCC ATT TAC CGT AAG TTA T-3’, Common Reverse: 5’-AAG GGA GCT GCA GTG GAG TA-3’. For collection of lung tissues, mice were sacrificed on P5 for creation of precision cut lung slices (PCLS). All animal work was approved by the Institutional Animal Care and Use Committee of Vanderbilt University (Nashville, TN) and was in compliance with the Public Health Services policy on humane care and use of laboratory animals. PCLS were created as described previously)(*4*). Briefly, lungs were inflated with low-melt temperature agarose and sliced on a vibratome to a thickness of 300-400 μm. Slices were washed in DMEM:F-12 medium with Penicillin / Streptomycin, then transferred to DMEM:F-12 without phenol red and with Penicillin / Streptomycin for imaging. Nuclei were stained with JaneliaFluor 646 at a concentration of 300 μM in the imaging medium (*16*). Tissues were maintained at 37 °C, 5% CO_2_, and atmospheric oxygen.

### Lung morphometry

To assess development in PCLS, tissues were fixed in 10% Phosphate buffered formalin at the time of creation (+0h) or 48h later. Fixed tissues were embedded in paraffin and thin sections were stained with hematoxylin and eosin. Airspace volume density was assessed as previously described (*4*), with observers blinded to experimental conditions.

### Light sheet imaging with scanned oblique plane illumination (SOPi) microscopy

Volumetric timelapse imaging was performed via two versions of SOPi microscope platforms, (the second system being a modified version of the first), which were built around the original design by Kumar et al. (*2, 17*). In brief, the OPM (oblique plane microscopy) class of light sheet platforms excite specimens at an oblique angle of incidence relative to the front lens of a single objective at the sample, serving both excitation and detection functions. By sweeping the angle of incidence of an offset, cylindrically-focused, excitation beam at the back focal plane of this objective, pure translation of an oblique planar (“light sheet”) excitation profile can be achieved at the sample. Thus, this approach maintains the low overall sample irradiance and high-speed capabilities inherent to many light sheet designs, while dramatically increasing sample flexibility and mounting practicality when compared to approaches requiring two (or more) orthogonally-oriented objectives (at the sample). While the emission side remote focus arrangement of OPM systems can be less efficient with lower overall system NA, the tradeoff of a) increased sample flexibility, b) high-speed volumetric imaging without the need to mechanically step the sample, and c) lower overall system complexity, results in a more practical solution for many non-conventional samples (such as precision cut lung slices), otherwise less amenable to alternative solutions.

We have made several updates to increase resolving power, reduce aberrations, and improve overall system sensitivity over the first SOPi build. First, in an effort to reduce aberrations associated with utilization of a standard achromat behind the primary imaging objective (MO1), we replaced this lens with a more highly corrected tube lens of the same focal length (ThorLabs, TTL-200MP). Second, the re-imaging objectives in the 2nd and 3rd microscope arm (MO2 and MO3) were replaced with the primary MO1 imaging objective model (Olympus 20x NA 1.0 WI, XLUMPLFLN20XW). This modification results in a higher effective NA of the system, thus increasing resolving power over the original design by almost twofold and reducing aberrations associated with mismatch between 1^st^ and 2^nd^ scope arms. In order to accommodate these updated re-imaging optics, a custom water chamber was fabricated for use between MO2 and MO3 objectives. Next, the sensor was also swapped for an Orca Fusion-BT sCMOS (Hamamatsu). A longer (f=400) focal length lens was installed prior to the updated sensor to increase the magnification and utilize more of the sensor array while maintaining Nyquist sampling, given the smaller pixel size (6.5 μm vs. 11 μm) vs. previous sensor. Excitation of fluorescence was accomplished by collimating the output of a single-mode fiber-based beam combiner (Galaxy, Coherent, Inc.) coupling 488, 561, and 640 -nm OBIS CW lasers (Coherent, Inc). Optical scanning of the light sheet was enabled by a large beam diameter, single axis, galvanometer (ThorLabs, Inc.). Sample placement and multi-point imaging (x,y,z) over time was possible via an automated (x,y) scanning stage and dual (z) linear stages (Applied Scientific Instrumentation). “Multi-point” imaging in this case indicates separate sample locations that were each optically scanned for volumetric imaging over the multi-day imaging runs (and not indicating a stage-scanned OPM approach, for example). This was both utilized for discrete location sampling as well as in image stitching applications to increase field-of-view. Live samples were environmentally controlled via stage top incubator regulating temperature (37°C), humidity (saturated, non-condensing), and 5% CO_2_ (Tokai Hit Co. LtD, Japan). Emission filtering for multichannel experimentation was accomplished via a triggerable emission filter wheel (Finger Lakes Instrumentation) and 525/50, 593/46, 615/20, 679/41 and 527/645 -nm emission filters (Semrock/Idex). Hardware triggering/timing between sensor, galvanometer, lasers, and filter wheel was coordinated via multifunction I/O board (PCIe-6353, National Instruments). Dataset and image acquisition was managed via NIS-Elements software (Nikon Instruments, Inc.) and Z8G4 workstation (HP, Inc) configured with 2x Intel 6244 CPU’s, 196GB RAM, solid state memory, and Quadro RTX-6000 GPU due to significantly increased computational demand incurred by size and nature of datasets. Such data represent volumetric imaging (~779 images/stack, with 0.45 μm/step) in multiple channels, at multiple stage coordinates, over several days of imaging. In total, PCLS were observed over 1,500 hours of imaging.

### Image acquisition and analysis

A multithreaded python-based image processing pipeline was used to efficiently process the large scale SOPi imaging data. Raw data from the microscope was first denoised in Nikon NIS-Elements, and then de-skewed using an affine transform using the SciPy python package (*18*) and re-saved in the OME-NGFF file format (*19*). Small drifts over the imaging time course were corrected by calculating the phase cross correlation between timepoints using the algorithm implemented in the Scikit-Image package (*20*) followed by adjusting the image location. 3D fluorescence stills and videos were created using the Napari image viewer (*21*) or Amira (ThermoFisher Scientific). Surfaces and cellular segmentations were determined by local thresholding with a neighborhood of one third of the imaging width (x axis) to account for the gaussian nature of the light sheet, followed by watershed segmentation to separate nearby discreet cells. Surfaces were determined using the marching cubes algorithm implemented in Scikit-Image. Nuclei were segmentized using Cellpose (*22*), and individual nuclei (or cells) are tracked through time with Bayesian Tracker (btrack) (*23*).

### Immunofluorescence

Immunofluorescence was performed as described previously (*4*). Briefly, 5 μm thick formalin-fixed, paraffin-embedded tissue samples on slides were deparaffinized and blocked with SeaBlock (ThermoFisher). Samples were stained with primary antibodies against GFP (1:100, ab13970 Abcam) and Ki67 (1:100, MA5-14520 ThermoFisher) followed by nuclear staining with DAPI. Images were acquired on a Keyence BZX-800. Quantification was performed using HALO software on images where the Ki67 signal was segmented to be counted only in nuclear areas.

### Computational model

Quantitative analysis of the alveologenesis data was performed on coordinates (*t*,*x*,*y*,*z*) of the inner boundaries of the alveolus bed. This data was separated by distinct timepoints resulting in *k* partitions of the data, each partition representing a mathematical snapshot of the shape of the alveolus boundaries at a particular time. Each snapshot (*t* = *T*) is analyzed by choosing a particular fixed value *z* = *Z* and studying the set of planar points: (*T,x*,*y*,*Z*). Starting with the *t* = 0 snapshot, closed, non-intersecting sets of points were identified. The set with the largest interior area is chosen as the alveolus of interest. At each instant of time, the set of points is approximated by a polygon at *z* = *Z*. The area of its interior and the perimeter length was calculated for the polygon. A program was written in Mathematica to perform these calculations, including fitting the set of points with a polygon. The sets {(*t_i_*, *A_i_*), *I* = 1, 2…, k} and {(*t_i_*, *P_i_*), *I* = 1, 2, ….k} were generated which describes the evolution of area and perimeter length over time. Another measure of growth/decay is the area divided by the perimeter length, the area-to-perimeter ratio is an indicator of polygon shape complexity and is the opposite of compactness.

---

**Figure S1:**
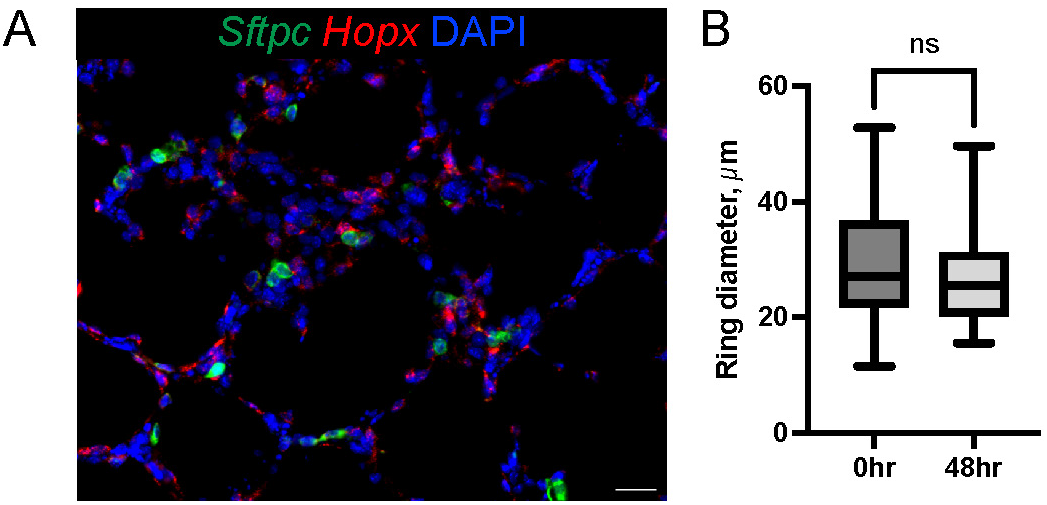
Characterization of PCLS. **A)** PCLS from P5 mouse after 48 hrs in culture. RNA *in situ* hybridization with probes for AT2 cell hallmark gene *Sftpc* (green) and AT1 cell hallmark gene *Hopx* (red) with DNA stained with DAPI (blue); scale bar = 25 μm. **B)** Measurement of alveolar mesenchymal ring diameter over 48 hr imaging in P5 PCLS from mT/mG;Pdgfra-Cre mouse, *p* = 0.34.

**Figure S2.**
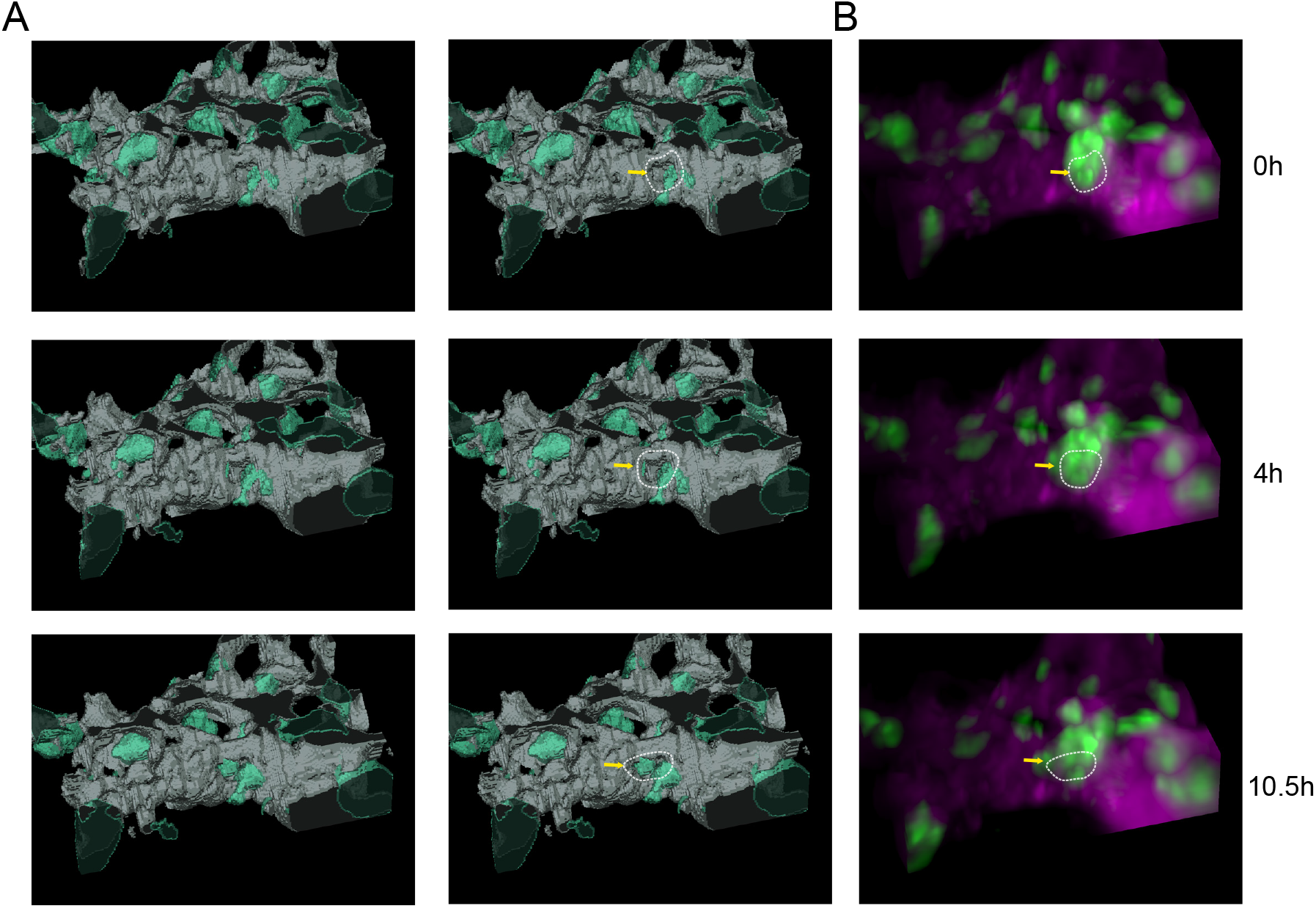
Epithelial outgrowths occur immediately adjacent to ring structures. **(A)** PCLS from mT/mG Nkx2-1-Cre mice were volumetrically imaged and displayed as a surface rendering where tdTomato positive cells are in gray and GFP positive cells are displayed in cyan. The dotted line indicates a ring structure immediately adjacent to an area of epithelial outgrowth indicated by the yellow arrow. **(B)** The source fluorescence image used for surface rendering where tdTomato positive cells are in magenta and GFP positive cells are green. The yellow arrow indicates an area of epithelial outgrowth.

**Figure S3:**
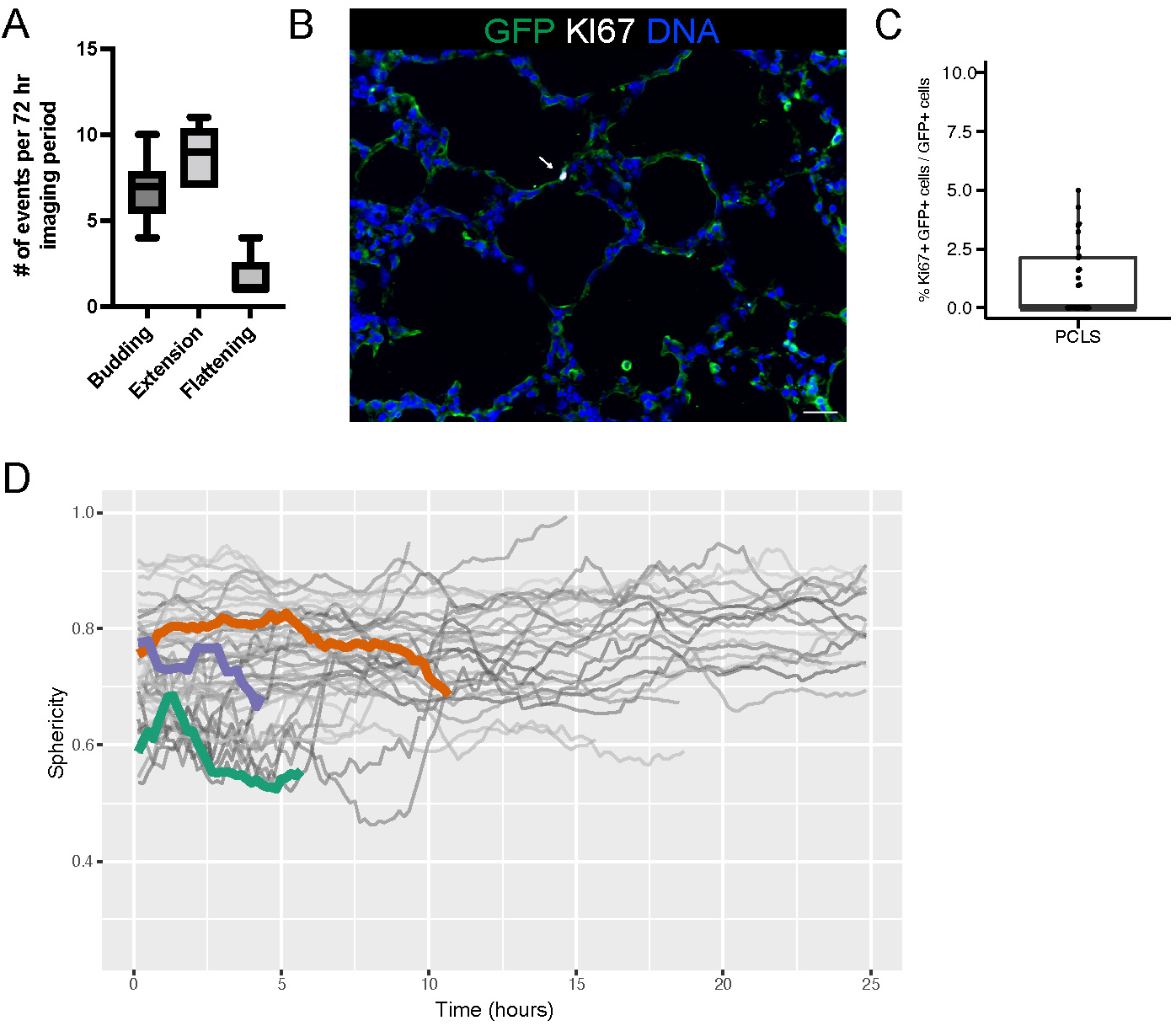
Alveologenesis at P5 is characterized by epithelial movement and characteristic shape changes, but limited replication. **(A)** Quantification of epithelial movements of Nkx2-1 labeled cells across 72 hr imaging period. **(B)** PCLS from a P5 mT/mG;Nkx2-1-Cre mouse was immunostained for GFP (green) and KI67 (white). DNA was stained with DAPI (blue). Arrow indicates a Ki67+, GFP+ cell. **(C)** The frequency of Ki67+ GFP+ cells was quantified using HALO software. **(D)** The sphericity was calculated from individually tracked GFP+ cells from a PCLS created from mT/mG;Nkx2-1-Cre mouse that was imaged over time. Gray lines indicate cells that did not flatten, while colored lines indicate cells that had a sphericity reduction of >= 0.1 over the course of imaging.

## Acknowledgments

Both custom scanned oblique planar illumination scopes (SOPi and MP-SOPi) were built by the BioMIID program within the Vanderbilt Biophotonics Center where images were also acquired, and resources utilized for visualization and analysis. Image acquisition was additionally performed in part through the use of the Vanderbilt Cell Imaging Shared Resource (supported by NIH grants CA68485, DK20593, DK58404, DK59637 and EY08126).

## Funding

National Institutes of Health grant T32HL094296 (NMN)

National Institutes of Health grant K08HL143051 (JS)

National Institutes of Health grant K08HL130595 (JAK)

National Institutes of Health grant R01HL153246 (JAK)

National Institutes of Health grant R01HL145372 (JAK)

National Institutes of Health grant P01HL092470 (TSB)

National Institutes of Health grant R01HL154287 (EJP)

National Institutes of Health grant R03HL154287 (EJP)

National Institutes of Health grant K08HL133484 (JTB)

National Institutes of Health grant R01HL157373 (JTB)

The Francis Family Foundation (JAK and JMSS)

The Chan Zuckerberg Initiative Imaging Scientists Program (BM)

Vanderbilt University Trans-Institutional Programs (TIPs) Award (BM, JK, AMJ)

## Author contributions

Conceptualization: NMN, BM, JMSS.

Data curation: NMN, BM, JMSS.

Formal analysis: NMN, YS, JMSS.

Funding acquisition: JMSS, BM.

Investigation: NMN, PC, YS, ANH, MR, CSJ, CB, MSS

Methodology: EJP, JTB, JMSS, BM, NMN, NM, AMJ, JK

Project Administration: CSJ, PG.

Software: NMN, BM, PC.

Validation: NMN, EJP, JTB, JAK, TSB, SHG, JMSS, ANH, MR.

Visualization: NMN, BM, YS.

Writing (original draft): NMN, JMSS.

Writing (review and editing): NMN, BM, YS, PC, EJP, DBF, WZ, CVEW, JTB, SHG, TSB, JAK, JMSS.

Supervision: JMSS, BM.

## Data and materials availability

All imaging data is available upon request. The code used for transforming and processing the SOPi imaging data is available at: https://github.com/SucreLab/SOPi_Alveologenesis

